# The Joint Action of Bacteriophage and Antibiotics in the Treatment of *Staphylococcus aureus* infections in *Galleria mellonella*

**DOI:** 10.64898/2026.04.13.718207

**Authors:** Brandon A. Berryhill, Teresa Gil-Gil

## Abstract

Given the global rise of antibiotic resistance, there has been a resurgence of interest in bacteriophage therapy, typically administered concomitantly with antibiotics and currently used as a last resort treatment. In this study, we use the *Galleria mellonella* model to investigate the treatment outcomes and dynamics of joint therapy against a toxigenic and pathogenic strain of *Staphylococcus aureus*. While our previous research demonstrated that single-agent therapy, whether using bactericidal or bacteriostatic antibiotics or lytic phage, could suppress infections below a critical threshold, it rarely achieved complete bacterial eradication. Here, we show that the coadministration of a phage PYO^Sa^ with antibiotics generally enhances clearance, regardless of the antibiotic class. Joint therapy with daptomycin resulted in the complete clearance of infecting bacteria in the majority of larvae. Notably, even when combined with ampicillin, to which the bacteria are highly resistant, approximately half of the larvae achieved infection clearance. Taken together, these results demonstrate that joint therapy with phage and antibiotics enhances clearance beyond what either agent achieves alone, while underscoring that treatment timing and drug-specific pharmacodynamics remain critical determinants of therapeutic outcome.

**Significance Statement:** The global rise of antibiotic resistance has renewed interest in bacteriophage therapy, which is almost universally administered concomitantly with antibiotics in clinical practice. Using *Galleria mellonella* (the wax moth larvae), which possess an innate immune response functionally similar to that of mammals, we demonstrate that coadministration of bacteriophages and antibiotics significantly enhances infection clearance compared with single-agent therapies. Critically, this joint action of antibiotics and phage can achieve bacterial eradication even when employing antibiotics to which the bacteria are resistant. We also find that therapeutic efficacy is sensitive to treatment timing and the specific pharmacodynamics of each drug. These factors are not captured by standard *in vitro* assessments of antimicrobial activity. Together, these results motivate further quantitative study of clinically relevant dosing regimens to determine the impact of host-pathogen-drug interactions on treatment outcome.

## Introduction

Due to the increasing frequency of antibiotic-resistant bacterial infections, there has been a renewed interest in the use of bacteriophage (phage) as a therapeutic (1). Currently, phage therapy is primarily used as a last-resort treatment (such as under Emergency Investigational New Drug authorizations in the USA) and is almost always used concomitantly with antibiotics (2, 3). Thus, there is a need to consider how these antimicrobial agents act in concert, as there have been disparate outcomes in phage therapy since its modern revival (4).

Previously, we have used the *Galleria mellonella* system to investigate the population and evolutionary dynamics of bacteria under single-agent therapy (both with antibiotics of different classes and phage) (5). While there are numerous considerations for using phage and antibiotics together, primarily the timing of administration, we first consider the simplest case: coadministration. In this brief report, we investigate the treatment outcomes of Galleria infected with *Staphylococcus aureus* MN8 treated with a highly lytic phage, PYO^Sa^, and with either a bactericidal or bacteriostatic antibiotic or a drug to which the infecting bacteria are resistant (6, 7). All four agents were investigated for their individual action in this system in our previous study. Here we demonstrate that the coadministration of both agents generally leads to lower bacterial loads recovered at both time points examined (4 and 24 hours), regardless of which ntibiotic was used. Our results raise the need to further investigate the role that different dosing regimens would have on treatment outcome to optimize therapeutic protocols which employ both antibiotics and phage.

## Results

In our investigation of the joint action treatment of antibiotics and the phage PYO^Sa^, we consider three different antibiotics: daptomycin (a traditionally bactericidal drug that is a first-line treatment for severe infections (8)), linezolid (a drug that is traditionally bacteriostatic for *Staphylococcus* and a second-line treatment (9, 10)), and ampicillin (a drug to which *S. aureus* MN8 is resistant (11)). All experiments used a single high-density inoculum (10^8^ CFU/larva), and three treatment conditions (immediate treatment sacrifice at 4h, immediate treatment sacrifice at 24 h, and treatment delayed until phenotypic markers of sickness with sacrifice at 24h). In Figure 1, daptomycin nearly completely abrogates morbidity and mortality (Supplemental Table 1). Despite the extremely high inoculum density, the average bacterial load sampled was below 10^1^ CFU/larva, with a majority (44/60; across the three treatment conditions) of the larvae being completely cleared of the infecting bacteria. However, despite having the lowest average bacterial density recovered, the delayed treatment has a higher degree of morbidity and mortality than the immediate treatment at either 4 or 24 hours. Linezolid, on the other hand, exhibits very different treatment outcomes and dynamics with immediate treatment (Figure 2); there is a higher degree of morbidity and mortality, as well as a higher bacterial load, with only three out of 40 larvae (across the two immediate treatment conditions) cleared of the infection (Supplemental Table 2). Linezolid likely impairs phage amplification, consistent with the lower phage densities recovered in the linezolid conditions (Figure 2B). The delayed treatments are similar between linezolid and daptomycin. Treating the larvae with an antibiotic to which the infecting bacteria are resistant and the phage (Figure 3) results in similar degrees of morbidity and mortality as for daptomycin (Supplemental Table 3), including approximately half of the larvae being completely cleared of the infection; however, the means of the bacterial densities recovered are similar to linezolid with immediate treatment.

**Figure 1.**
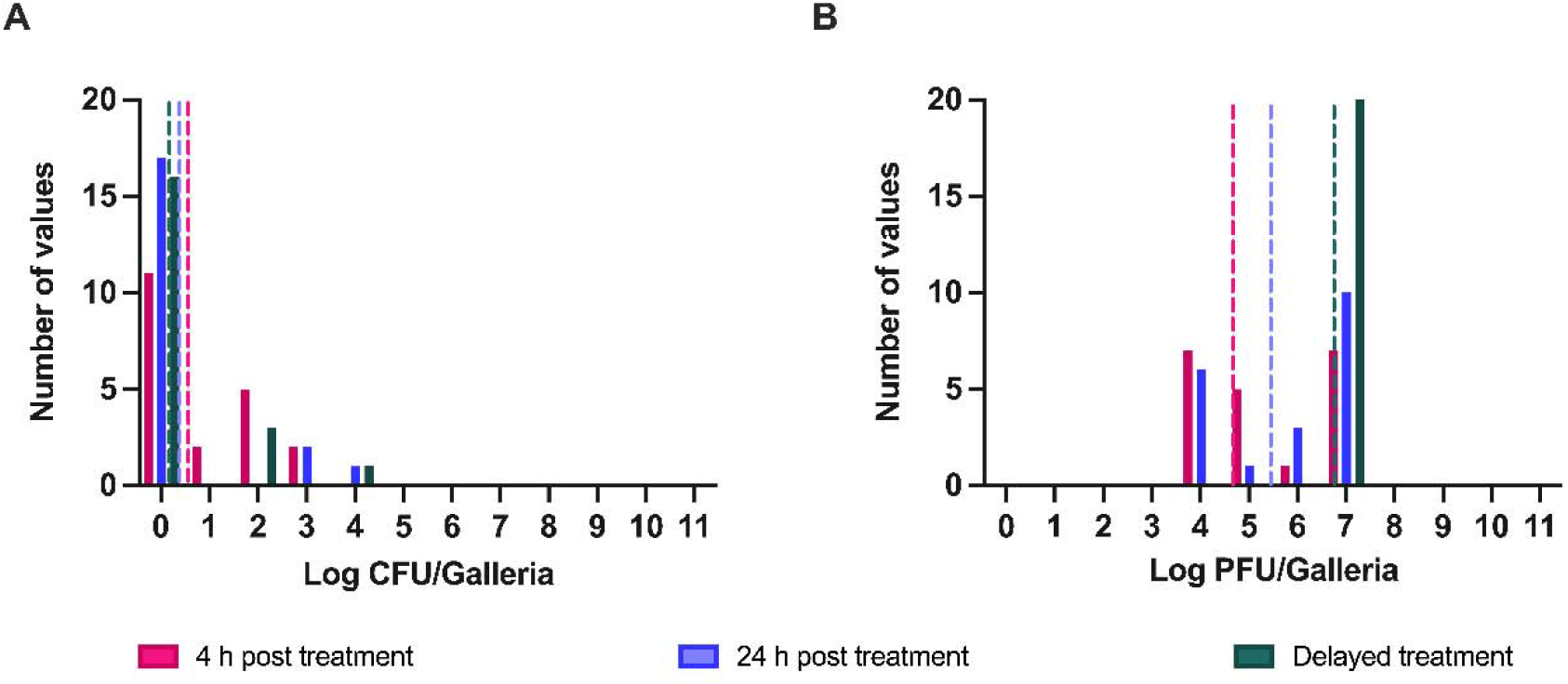
Dynamics of infection with daptomycin and phage treatment. Larvae were infected with high densities (10^8^ CFU/larva) of *S. aureus* MN8 and were sacrificed after 4 (pink) or 24 (blue and green) hours post-infection. Treatment occurred either immediately with 10^8^ PFU/larva and the antibiotic (pink and blue) or once symptoms of infection appeared (green). Shown are the number of larvae within each log_10_ CFU (A) or the number of larvae within each log_10_ PFU (B) for each respective experiment. The mean of the CFU or PFU is presented as a dotted line. For each condition, N=20 individual larvae were inoculated. Survival and morbidity data are reported in Supplemental Table 1.

**Figure 2.**
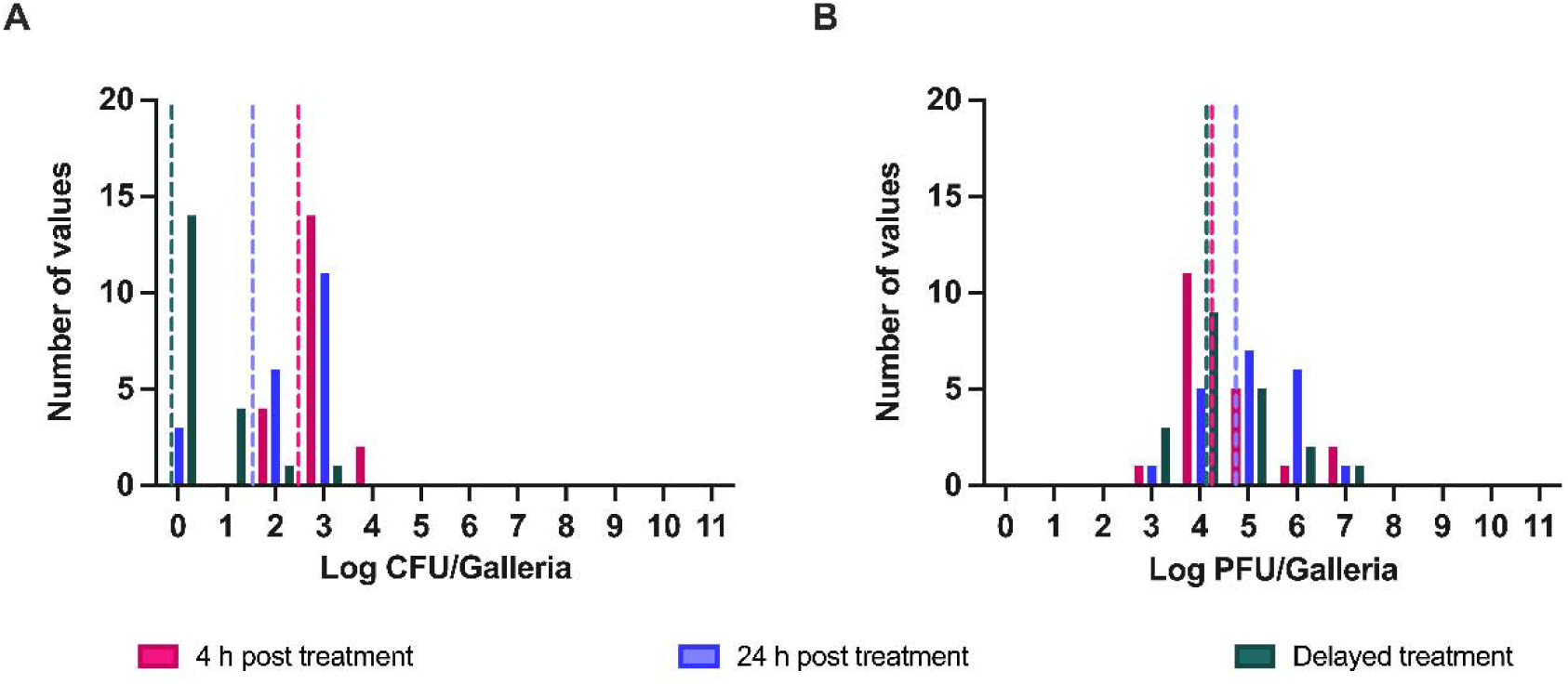
Dynamics of infection with linezolid and phage treatment. Larvae were infected with high densities (10^8^ CFU/larva) of *S. aureus* MN8 and were sacrificed after 4 (pink) or 24 (blue and green) hours post-infection. Treatment occurred either immediately with 10^8^ PFU/larva and the antibiotic (pink and blue) or once symptoms of infection appeared (green). Shown are the number of larvae within each log_10_ CFU (A) or the number of larvae within each log_10_ PFU (B) for each respective experiment. The mean of the CFU or PFU is presented as a dotted line. For each condition, N=20 individual larvae were inoculated. Survival and morbidity data are reported in Supplemental Table 2.

**Figure 3.**
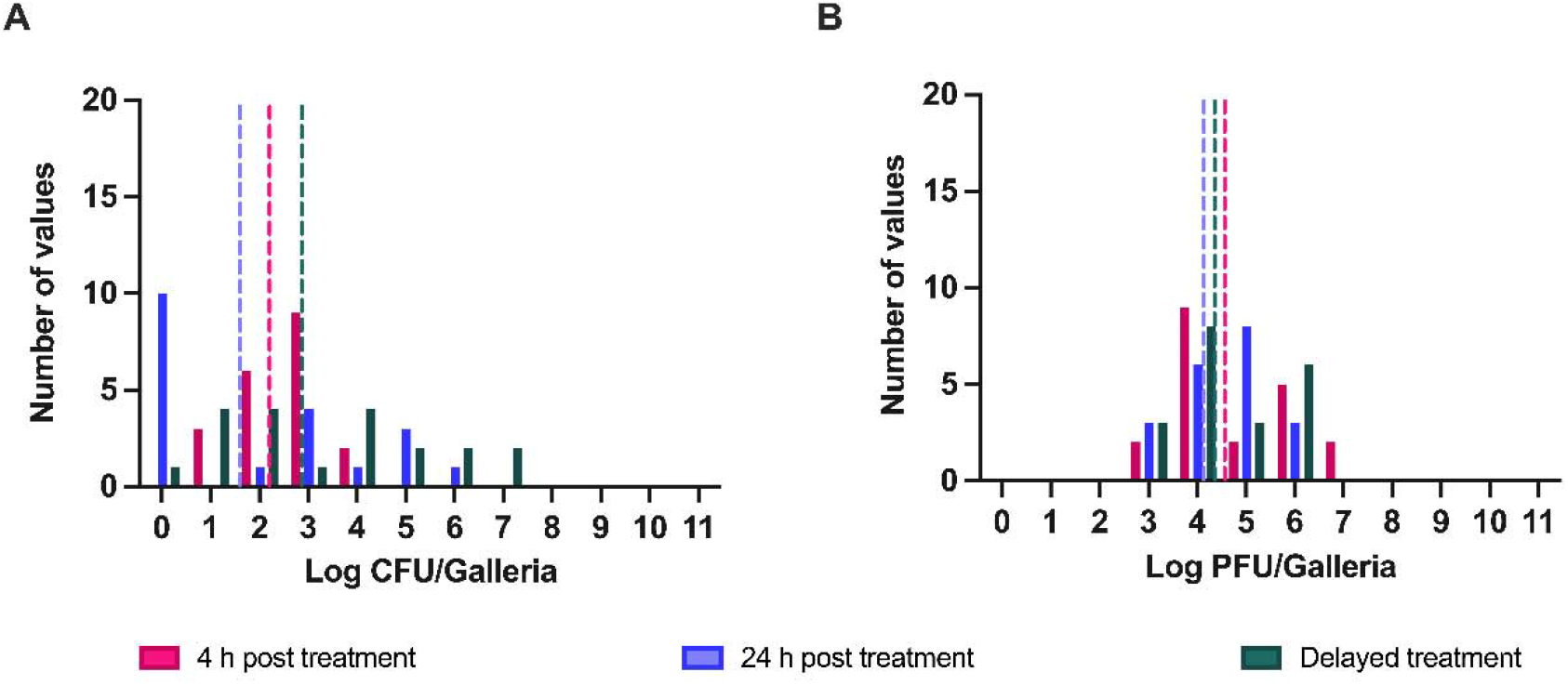
Dynamics of infection with ampicillin and phage treatment. Larvae were infected with high densities (10^8^ CFU/larva) of *S. aureus* MN8 and were sacrificed after 4 (pink) or 24 (blue and green) hours post-infection. Treatment occurred either immediately with 10^8^ PFU/larva and the antibiotic (pink and blue) or once symptoms of infection appeared (green). Shown are the number of larvae within each log_10_ CFU (A) or the number of larvae within each log_10_ PFU (B) for each respective experiment. The mean of the CFU or PFU is presented as a dotted line. For each condition, N=20 individual larvae were inoculated. Survival and morbidity data are reported in Supplemental Table 3.

Survival with delayed ampicillin and phage treatment was notably higher than with delayed linezolid and phage, though lower than delayed daptomycin and phage, with only one larva out of 20 being cleared. In terms of phage densities recovered from all larvae, the daptomycin treatment groups consistently had higher densities of phage recovered, as unlike linezolid, there is no mechanistic reason to expect antagonism.

## Discussion

The results of this study demonstrate that the simultaneous coadministration of bacteriophage and antibiotics generally enhances the clearance of *S. aureus* MN8 infections in *G. mellonella* compared to the individual action of these agents. While our previous work established that single-agent therapy, whether using bactericidal or bacteriostatic antibiotics, or lytic phage, could successfully control infections by reducing bacterial loads below a critical threshold of approximately 10^6^ CFU/larva, those treatments rarely achieved complete clearance (11). In contrast, joint therapy with daptomycin and the phage PYO^Sa^ resulted in the complete eradication of infecting bacteria in the majority of larvae. Even with ampicillin, a drug to which the bacteria are highly resistant, in combination with phage resulted in infection clearance in approximately half of the larvae. This suggests that the two agents, acting together with the host’s innate immune system, can overcome the consistent lack of clearance observed in single-treatment regimens.

*G. mellonella* is an exceptionally tractable *in vivo* system for testing quantitative hypotheses in infection biology (12-14). These larvae possess an innate immune response analogous to that of mammalian neutrophils, infections are highly replicable, and the phenotypic markers of morbidity allow treatment to be initiated once symptoms are present rather than at a fixed time (15). Our results demonstrate that this system is well-suited for studying the joint action of antimicrobial agents under conditions that reflect the clinical reality of infection.

The scope of the present study is deliberately restricted to the simplest case, coadministration, as a necessary first step before investigating more complex dosing regimens. It is also worth noting that, like all invertebrate models, *G. mellonella* lacks an adaptive immune system, and the contribution of adaptive immunity to the dynamics of joint therapy remains an open question. Given these limitations, our data show that delayed treatment administered once symptoms of infection are present, results in higher morbidity and mortality compared to immediate treatment, especially with linezolid or ampicillin combinations. This highlights that the efficacy of joint therapy with antibiotics and phage is sensitive to the state of the infection and the specific pharmacodynamics of the antibiotics used. These observations underscore the next phase of investigation.

Moving forward, there is a critical need to investigate more complex dosing regimens to optimize therapeutic protocols. Of particular interest would be investigating sequential vs. simultaneous dosing to test whether phage-first or antibiotic-first timing alters outcomes (16). Further, our results raise the question of how different timing intervals between antibiotic and phage administration might influence the ultimate trajectory of the infection. Future research should also explore the evolutionary dynamics of resistance during joint therapy, as our previous work showed using immunodeficient *Galleria* that the innate immune system is the primary factor preventing the ascent of resistant minority populations (11). By taking advantage of the *G. mellonella* system’s high replication capacity and its phenotypic markers of sickness, we can begin to design treatment regimens that more accurately reflect human clinical scenarios, where therapy is initiated only after the host is visibly ill.

## Materials and Methods

### Growth media and conditions

All experiments were conducted in Mueller-Hinton II (MHII) Broth (90922-500G) obtained from Millipore. All bacterial quantification was done on Lysogeny Broth (LB) agar (244510) plates obtained from BD. All experiments were conducted at 37 °C.

### Bacterial and bacteriophage strains

All experiments were performed with *S. aureus* MN8 obtained from Tim Read of Emory University. *S. aureus* MN8 was marked with streptomycin resistance to enable differential plating from the larva microbiota. The bacteriophage PYO^Sa^ was obtained from the Levin Laboratory’s bacteriophage collection. Bacteria and phage were prepared for injection and plated as in (Cite Worm 1).

### Antibiotics

Daptomycin (D2446) was obtained from Sigma-Aldrich. Streptomycin (S62000) was obtained from Research Products International. Ampicillin (A9518-25G) was obtained from Sigma-Aldrich. Linezolid (A3605500-25g) was obtained from AmBeed.

### *G*. *mellonella* preparation

*G. mellonella* larvae were obtained from Premium Crickets (Georgia, USA) and placed immediately at 4 °C for 48 hours as a cold shock. Larvae were then sorted such that only those that weighed between 180 and 260 mg.

### *G*. *mellonella* treatment

Larvae were treated by injection in the last right proleg. Antibiotic concentrations were analogous to those used for clinical treatment in humans and the amount of antibiotic determined by weight for a total final treatment amount of: LIN-0.002 mg per larva; DAP-0.002 mg per larva; AMP-0.04 mg per larva. Phage was delivered in the same way with a fixed concentration of 10^8^ PFU per larva.

## Acknowledgments

We would like to thank Alysha Ismail for her excellent laboratory management work and our respective mentors Dan I. Anderson and Bruce R. Levin as well as the rest of the Anderson Lab and Levin Lab for their support.

## Supplemental Materials

**Supplemental Table 1.** Daptomycin treatment survival.

| **Treatment** | **Healthy** | **Sick** | **Dead** |
| --- | --- | --- | --- |
| **Immediate Kill at**<br>**4 Hours** | 19 | 1 | 0 |
| **Immediate Kill at**<br>**24 Hours** | 19 | 0 | 1 |
| **Delayed**<br>**Treatment Kill at**<br>**24 Hours** | 12 | 6 | 2 |

**Supplemental Table 2.** Linezolid treatment survival.

| **Treatment** | **Healthy** | **Sick** | **Dead** |
| --- | --- | --- | --- |
| **Immediate Kill at**<br>**4 Hours** | 16 | 4 | 0 |
| **Immediate Kill at**<br>**24 Hours** | 5 | 10 | 5 |
| **Delayed**<br>**Treatment Kill at**<br>**24 Hours** | 14 | 2 | 4 |

**Supplemental Table 3.** Ampicillin treatment survival.

| **Treatment** | **Healthy** | **Sick** | **Dead** |
| --- | --- | --- | --- |
| **Immediate Kill at 4 Hours** | 16 | 4 | 0 |
| **Immediate Kill at 24 Hours** | 18 | 0 | 2 |
| **Delayed Treatment Kill at 24 Hours** | 8 | 7 | 5 |

